# OPTIMIZATION OF DRIED BLOOD SPOT ASSAYS FOR HEPATITIS B VIRUS SURFACE ANTIBODY QUANTIFICATION

**DOI:** 10.1101/2024.05.23.595500

**Authors:** Patience Motshosi, Bonolo B. Phinius, Mosimanegape Jongman, Kabo Baruti, Lynnette Bhebhe, Wonderful T. Choga, Sikhulile Moyo, Simani Gaseitsiwe, Motswedi Anderson

**Affiliations:** Botswana Harvard Health Partnership, Gaborone, Botswana; School of Allied Health Professions, Faculty of Health Sciences, University of Botswana Gaborone, Botswana; Department of Biological Sciences, University of Botswana, Gaborone, Botswana; Department of Immunology and Infectious Diseases, Harvard T.H. Chan School of Public Health, Boston, Massachusetts, USA; School of Health Systems and Public Health, University of Pretoria, South Africa; Division of Medical Virology, Faculty of Medicine and Health Sciences, Stellenbosch University, Tygerberg, South Africa; University of KwaZulu Natal

**Keywords:** dried blood spots (DBS), HBV, HBsAg, anti-HBs, in-house assay

## Abstract

Dried blood spot (DBS) cards can be used as an alternative sample collection method to plasma, however, there is no optimized elution protocol for DBS cards specifically for hepatitis B surface antibody (anti-HBs) testing. The study aimed to develop a DBS elution protocol for anti-HBs quantification. Our study sought to determine the ideal phosphate-buffered saline (PBS) buffer volume to use by comparing three PBS volumes (300uL, 450uL, and 500uL), and the optimal time to agitate DBS discs on a plate shaker (1hr, 2hrs, 3hrs, and 4hrs) to yield DBS anti-HBs concentrations that are comparable to corresponding plasma anti-HBs concentrations. The optimal DBS storage temperature (25°C, -20°C, and -80°C) was investigated to determine the ideal longterm storage temperature of the cards. Residual samples were used for optimization (2019-2021). A total of 50 DBS-plasma pairs was used throughout the study, with plasma anti-HBs concentrations being used as the golden standard to compare. The analysis of results was carried out by determining the p-values of the Wilcoxon sign rank. A two-way analysis of variance (ANOVA) was also performed to determine the impact of PBS elution volumes, elution time, and storage temperature on the anti-HBs concentration of DBS samples on STATA Version 15.0. No statistically significant difference between the DBS-plasma anti-HBs pairs was observed when using 450 or 500uL of PBS buffer and when samples were agitated for 3 hours (p=0.594, p=0.499 respectively). The optimal storage temperature for DBS cards was 25°C because the results showed no statistically significant difference between DBS-plasma anti-HBs titers (p=0.594). The two-way ANOVA analysis showed that elution volumes and time had no statistically significant impact on the DBS anti-HBs concentrations, p=0.948 and p=0.381 respectively. Storage temperature had a statistically significant impact on the DBS anti-HBs concentrations, p=0.002. The optimized DBS elution protocol for anti-HBs quantification will help monitor vaccine efficacy in infants due to the low sample volumes required compared to plasma and also can be used for anti-HBs testing in resource-limited areas around the country.

## INTRODUCTION

Hepatitis B is a liver infection that is caused by the hepatitis B virus (HBV), which leads to complications such as hepatocellular carcinoma (HCC), and cirrhosis [1]. It is a significant global health problem that is estimated to affect 296 million people globally who are chronically infected, leading to 820,000 deaths per year [1]. The prevalence of chronic HBV infection among adults is highest in the Western Pacific (6.2%) and African regions (6.1%) [1]. In Botswana, the HBV prevalence rate has been reported to range between 1.1 – 10.6% among different key populations [1-4].

There is an increasing focus on coming up with strategies aimed at achieving the World Health Organization (WHO) target to eliminate HBV as a public health concern by 2030, defined as reducing HBV incidence by 90% and mortality by 65% [3]. The age at infection primarily determines the rate of progression from acute infection to chronic infection, which is approximately 90% in the perinatal period and 20-50% in children aged 1-5 years [3]. The HBV vaccine remains the mainstay of reducing HBV incidence, with a target of reduction in the prevalence of HBV surface antigen (HBsAg) in children to 0.1% by 2030 [3]. Consequently, HBV screening and monitoring of vaccine immune responses remain central to achieving this aim. The current method of monitoring response to the HBV vaccine involves the quantification of hepatitis B surface antibodies (anti-HBs) using Enzyme-linked Immunosorbent assay (ELISA), which relies on the use of plasma or serum as sample types [5]. A previous study used plasma samples from infants who were exposed to the human immunodeficiency virus (HIV) but uninfected to determine the HBV vaccine response in Botswana and insufficient volume limited testing of other HBV biomarkers [3]. More recently, another study utilized dried blood spot (DBS) samples to determine the immune response of infants with HIV and those without HIV in Botswana [6]. However, data such as optimal DBS storage temperatures were not provided by the study.

The challenge with plasma or serum samples is that they require freezer storage to maintain longterm sample integrity [7]. Furthermore, these sample matrices are unsuitable for use in monitoring vaccine response among infants due to the large sample volume required for collection [8]. Lowand middle-income countries lack basic storage facilities such as freezers due to their high cost and have an unreliable electricity supply to support such important sample storage equipment [9]. Therefore, middle-income countries like Botswana are forced to consider alternative sample collection methods, such as dried blood spots (DBS) to meet their needs. The advantages of using DBS for monitoring vaccine response is that it can be prepared with whole blood collected from a finger stick, causing the patient less discomfort, it does not require equipment such as centrifuge machines and freezers and desiccated samples can be stored for transport as non-hazardous material via postal services [9]. This makes the use of DBS cards ideal for monitoring HBV vaccine response in infants as well as in remote areas with limited healthcare resources.

Therefore the study aimed to use paired DBS-plasma samples to optimize a working DBS elution protocol for anti-HBs quantification to use DBS cards as an alternative sample collection method to plasma/serum.

## METHODS

### Study Participants

This was a retrospective cross-sectional study utilizing paired DBS cards and plasma samples (n=50) obtained from the CD4 laboratory at Botswana Harvard Health Partnership (BHP) on the 22^nd^ of March 2021 and all the authors had no access to information that could identify individual samples. These were residual samples anonymous (delinked) from the integrated patient management system (IPMS) with respect to the medical confidentiality of patients to only be used for optimization purposes, therefore, no consent was seeked as sample identities were hidden. The ethics review board of the Ministry of Health and Wellness, Botswana issued a research permit to perform the study at the Botswana Harvard Health Partnership, reference number: HPDME:13/18/1. Whole blood was collected on Whatman filter paper 903 (GE Healthcare, Piscataway, NJ) and stored at different temperatures (-80°C, -20°C, and 25°C) for 26 months before optimization.

### Optimization of DBS elution protocol

#### Optimization of PBS volumes

A set of 10 DBS cards stored at 25°C were punched into 2ml Eppendorf tubes and suspended in three varying elution volumes of PBS buffer (300μl, 450μl, and 500μl). The tubes were continuously agitated at 1600 revolutions per minute (rpm) on a plate shaker. The discs were removed, and the tubes were centrifuged at 10500 rpm for 2 minutes. The anti-HBs concentrations of the DBS samples were then tested using the Monolisa Anti-HBs PLUS kit (Bio-Rad, Hercules, CA, USA) with a 2.00 mIU/mL lower limit of detection (LoD) and 1,000 mIU/mL upper LoD, following the manufacturers’ instructions. The results were compared to their corresponding plasma anti-HBs concentrations to determine the best PBS buffer volume to use when eluting DBS cards.

#### Optimization of Elution time

A different set of 10 DBS cards was used to determine the best time to shake/agitate DBS discs after being suspended in 500μl of PBS buffer [6]. This set of 10 DBS discs was agitated on a shaker at 1600rpm for varying time intervals (1 hour, 2 hours, 3 hours, and 4 hours). The discs were removed, and the tubes were centrifuged at 10500 rpm for 2 minutes. The anti-HBs concentrations of the DBS samples were tested using the Monolisa Anti-HBs PLUS kit (Bio-Rad, Hercules, CA, USA) with a 2.00 mIU/mL lower limit of detection (LoD) and 1,000 mIU/mL upper LoD, following the manufacturers’ instructions. The results were compared to their corresponding plasma anti-HBs concentrations to determine the best elution time for DBS discs.

#### Optimization of storage temperatures

After determining the ideal PBS elution volume and time, anti-HBs quantification was performed on a set of 30 DBS-plasma pairs with each 10 DBS cards stored at a different temperature to determine the optimum storage conditions for DBS cards. The DBS discs were placed in 2ml Eppendorf tubes containing 500μl of PBS buffer and were agitated for 4 hours at 1600rpm on a shaker. These DBS cards were stored at three varying storage temperatures (-80°C, -20°C freezers, and 25°C). The discs were removed, and the tubes were centrifuged at 10500 rpm for 2 minutes. The anti-HBs concentrations of the DBS samples were tested using the Monolisa Anti-HBs PLUS kit (Bio-Rad, Hercules, CA, USA) with a 2.00 mIU/mL lower limit of detection (LoD) and 1,000 mIU/mL upper LoD, following the manufacturers’ instructions. The results were compared to their corresponding plasma anti-HBs concentrations to determine the best elution time for DBS discs.

### Data analysis

Optimum PBS elution volume, elution time, and storage temperature were determined by comparing anti-HBs results from DBS cards to their corresponding plasma aliquots. Statistical analysis was performed using STATA Version 15.0. The Wilcoxon signed-rank test was used to determine the p-values by comparing the anti-HBs concentrations of DBS samples to their corresponding plasma aliquots and two-way ANOVA was used to determine the statistically significant impact of elution volume, elution time, and storage temperature on the anti-HBs concentrations. P<0.05 showed a statistically significant difference. Box plots were used to show the distribution of data of the DBS and plasma pairs and help compare the distributions to each other.

### Work Flowchart

## RESULTS

The box plots below show the distribution of data for the DBS vs plasma anti-HBs concentrations.

The distribution of the data for the DBS samples and plasma aliquots is similar, the p-values are all greater than 0.05. The median lines of the plots are closer to each other (a.), indicating that eluting DBS discs with either 300, 450, or 500uL will not result in a significant difference between the DBS vs plasma anti-HBs concentrations. The closer the median lines are between the DBS and plasma box plot, the better the elution volume hence 450uL was the best elution volume (figure 2a). Due to the outliers (b.) the box plots become obscured making interpretation of the plots difficult. Box plot (a.) has no outliers while box plot (b.) has outliers.

**Figure 1:**
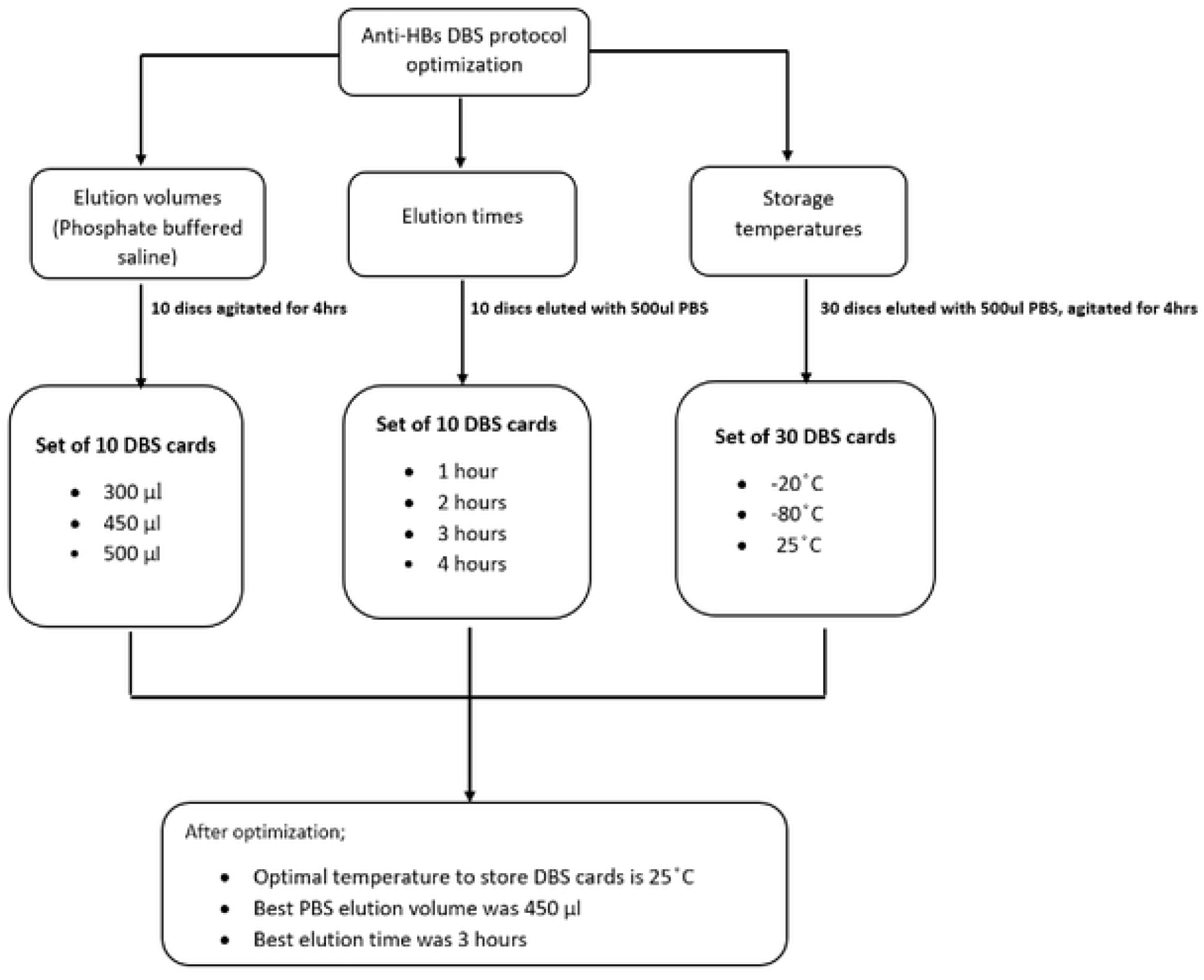
Optimization of DBS elution protocol for anti-HBs quantification

**Figure 2:**
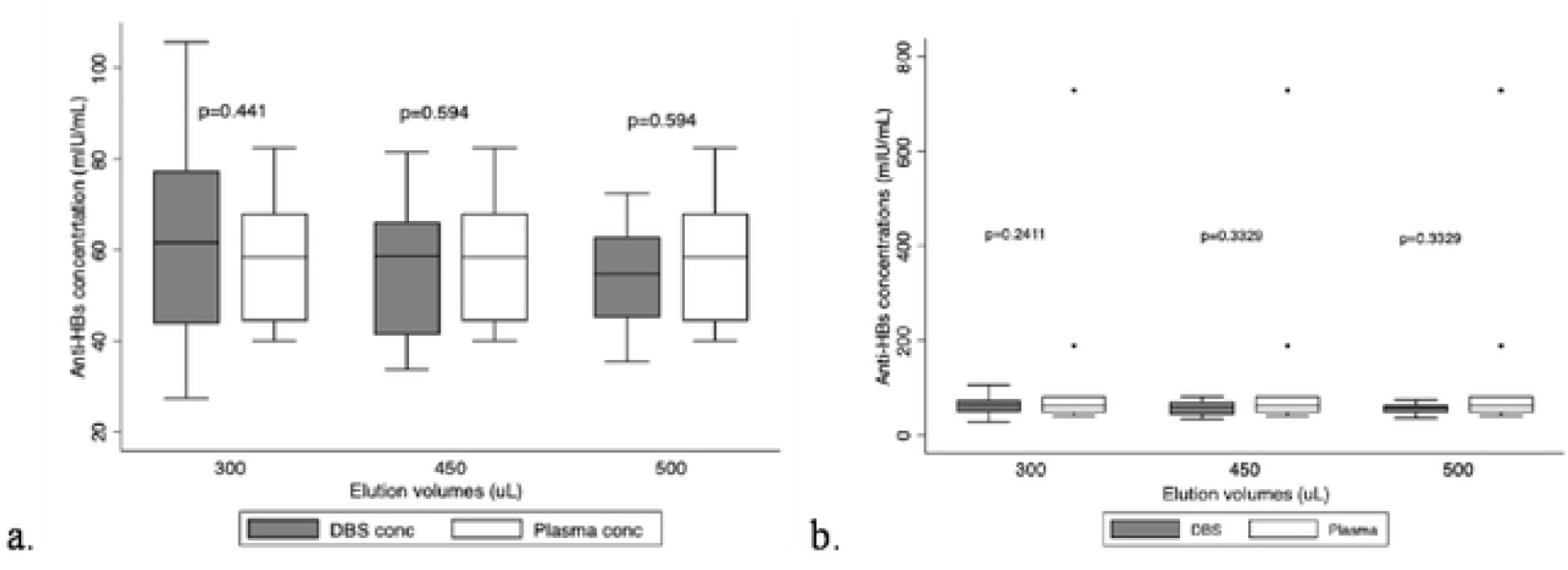
Box plot with and without outliers comparing the different PBS elution volumes

**Figure 3:**
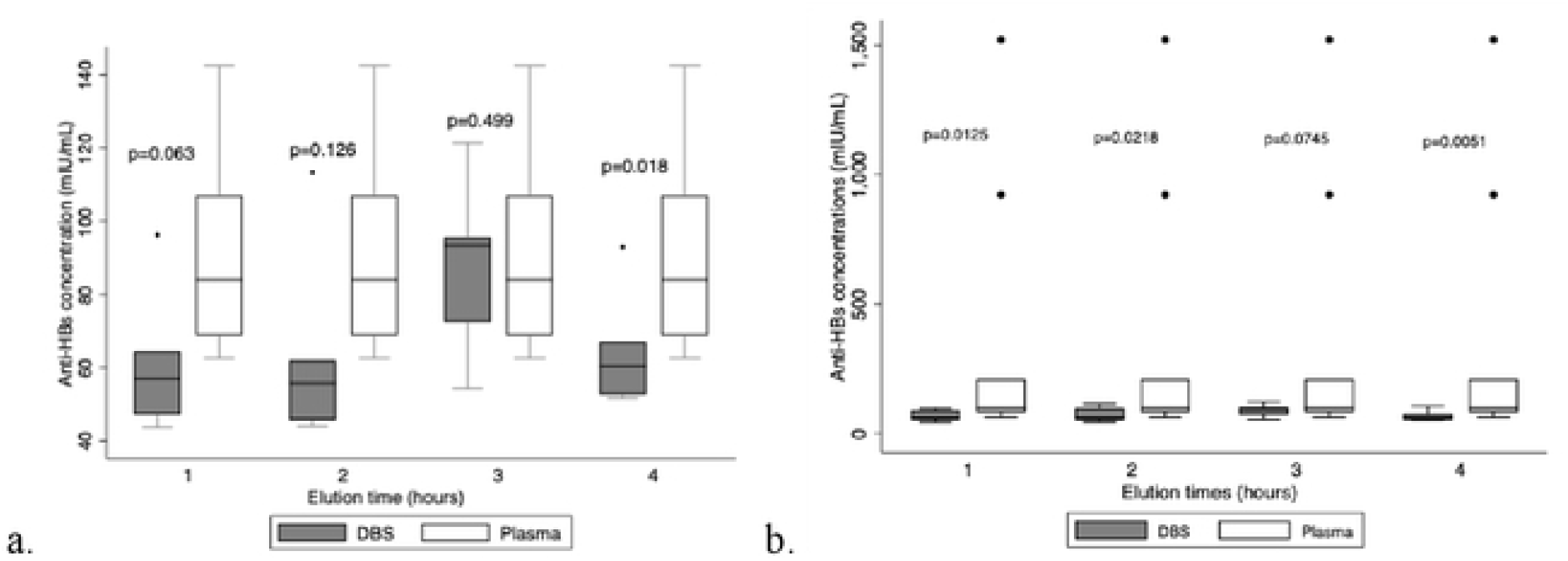
Box plot with and without outliers comparing the different times DBS discs were agitated

**Figure 4:**
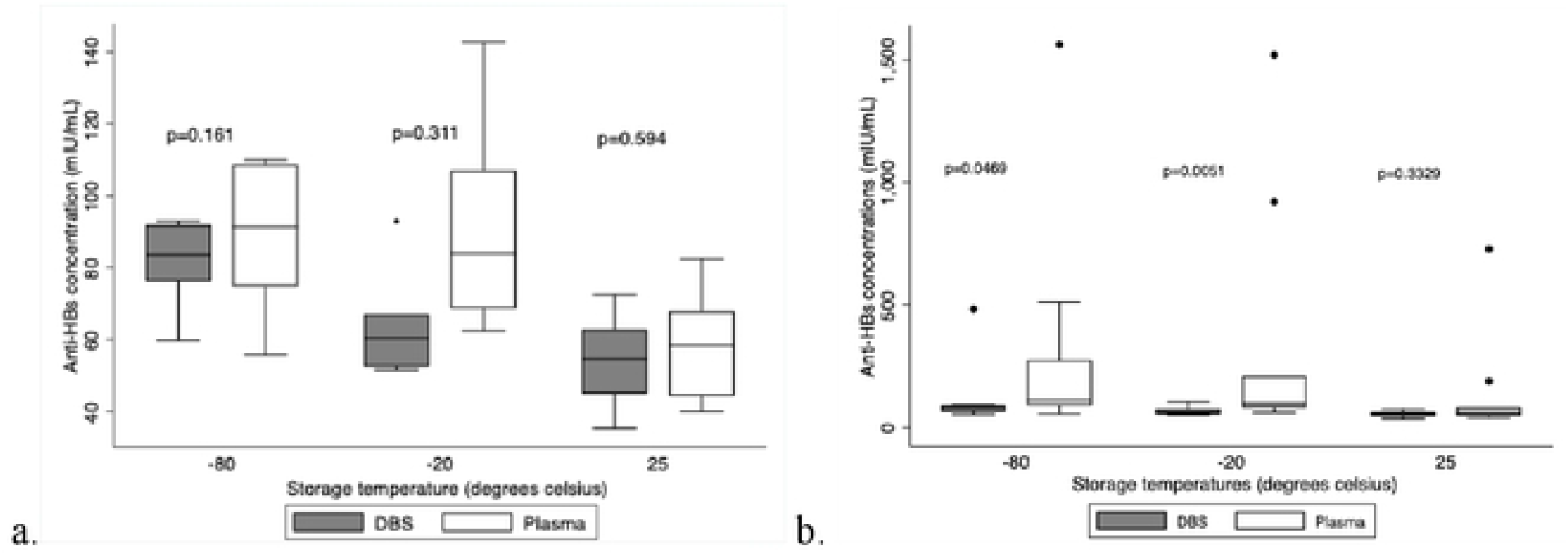
Box plots with and without outliers comparing anti-HBs concentrations for DBS cards stored at different temperatures

The distribution of the DBS and plasma plots, along with the p-value of 0.499 (a.), indicates that agitating DBS discs on a plate shaker is best done for three hours. This is in contrast to the other three elution times, which show a statistically significant difference between the concentration of anti-HBs in plasma and DBS discs, based on upper and lower quartiles of the DBS vs. plasma plots that are considerably different from one another. The distribution of the data at 1, 2, and 4 hours indicates that the plasma points are not as widely distributed as those of the DBS, indicating an inconsistency in the data distribution (a.). In contrast to the three other time intervals that revealed a statistically significant difference in the concentrations, the p-value at 3 hours also reveals no statistically significant difference between the DBS vs. plasma anti-HBs concentration

For long-term storage, DBS cards can be kept at 25°C, according to the box plot and p-values. When comparing the anti-HBs concentration of DBS to those of plasma, the p-value of 0.594 indicates that there is no significant difference, and the closeness of the median lines of the box plots of plasma and DBS at 25°C indicates that outcomes are similar (a.). The data distribution for the DBS vs. plasma plots at 25°C also coincides with the p-value, indicating that this was the best temperature to store DBS cards for an extended period of time. Box plot (a.) has no outliers while box plot (b.) has outliers, this shows how the outliers affect the interpretation of the data.

A two-way ANOVA was performed to analyze the effect of PBS elution volumes, elution time, and storage temperature on the anti-HBs concentration of DBS samples. It revealed that there was a statistically significant interaction between the effects of PBS elution volume, elution time, and storage temperature on the anti-HBs concentrations of DBS samples (F=2.30, p=0.034).

**Table 1:** p-values of two-way analysis of variance (ANOVA)

| Variables | p-values | F | Sums of sq. | df | MS |
| --- | --- | --- | --- | --- | --- |
| Volume | 0.948 | 0.05 | 209.95349 | 2 | 104.97675 |
| Time | 0.381 | 1.04 | 6041.4985 | 3 | 2013.8328 |
| Storage temperature | 0.002 | 6.49 | 25268.302 | 2 | 12634.151 |
| Volume, Time & Storage Temperature | 0.034 | 2.30 | 31260.555 | 7 | 4465.7935 |
| Volume & Time | 0.737 | 0.55 | 5992.2533 | 5 | 1198.4507 |
| Volume & Storage temperature | 0.016 | 3.24 | 25219.056 | 4 | 6304.7641 |
| Time & Storage temperature | 0.009 | 3.26 | 31050.601 | 5 | 6210.1203 |

## DISCUSSION

Our study sought to discover the ideal conditions for using DBS cards as an alternative sample collection method for the quantification of anti-HBs by optimizing three different variables. The study took into account the ideal PBS elution volume, the amount of time needed to successfully elute the DBS discs, and the optimal storage temperature to store DBS cards. DBS cards that were stored at 25°C, eluted using 450uL PBS buffer, and then placed on a plate shaker for 3 hours revealed excellent agreement between DBS samples and their respective plasma aliquots.

The study findings in terms of the effect of elution volume on anti-HBs concentration are in line with previous studies [6], as it has revealed that eluting DBS cards with 450ul will yield concordant anti-HBs concentrations between DBS-plasma pairs. This research has also shown that PBS, a commonly used elution buffer, aids in maintaining the stability of antibodies during sample processing [10]. It was discovered that 450ul was the ideal elution volume, as it eluted most of the disc and was enough to cover and saturate the DBS discs. According to [6], this volume provides a balance between optimal antibody recovery and reducing potential differences in elution effectiveness due to over-flooding the DBS discs and over-saturation of the discs. The range of 450ul to 500ul of PBS buffer also considers factors such as the DBS disc’s saturation capacity, which helps avoid having a dry or overly soaked DBS disc, to guarantee an effective elution procedure [11].

The literature has also demonstrated how important it is to optimize a DBS elution conditions for anti-HBs quantification. Considering the widespread use of DBS cards in both laboratory and field settings, it makes sense logically to use DBS cards for their practicability in resource-limited areas and as an efficient means of enabling sample collection and extraction [11]. A study by [6] has demonstrated that eluting DBS discs for 4 hours will yield concordant anti-HBs concentrations between DBS-plasma pairs. This is contradictory to our study as we have shown that eluting DBS discs for longer than 3 hours yields discordant anti-HBs concentrations between DBS-plasma pairs. Our suggested elution time of 3 hours for shaking has practical implications as it reduces the possibility of antibodies becoming unstable due to ongoing friction [6]. This finding is particularly significant for resource-limited settings where it may be challenging to maintain laboratory conditions appropriate for the elution procedure, ensuring a practical and efficient elution process for anti-HBs testing [6].

Different regions have differing average temperatures, therefore the need to have a practical and standard storage temperature that can be applied in all settings is important. 25°C is a practical storage temperature especially taking into account that this optimization study was targeting to optimize the use of DBS cards in resource-limited settings. To maintain the integrity of anti-HBs throughout storage and ensure the precision of serological testing carried out on DBS samples an optimal storage temperature needs to be established [11]. The results from this study add to the growing body of knowledge regarding the use of DBS cards as an alternative sample collection method and its potential to enhance diagnostic precision, especially in low and middle-income countries like Botswana where access to advanced laboratory infrastructure is scarce in many rural areas. We have demonstrated and provided evidence that the ideal temperature for storing DBS cards for extended periods of time is 25°C. This temperature has preserved the antibodies and kept them stable for longer than 2 years. This is contrary to what another study has discussed, [8] showed that DBS for anti-HBs quantification is a reliable alternative to plasma specimens when stored at -20°C and that antibodies will only remain stable for up to 183 days at room temperature (20°C-25°C). Therefore, moving forward, it may be useful to determine the stability of antibodies on DBS cards at various temperatures when stored for varying periods. Our study is currently the only study in Botswana to have used DBS cards stored for longer than 2 years. This brings a different aspect that shows the practicability of DBS cards, particularly in resource-limited areas in Botswana. This will potentially facilitate the testing of vaccine efficacy, especially in infants and in less developed areas of the country.

An analysis of simple main effects using two-way ANOVA indicated that the PBS elution volume had no statistically significant impact on the anti-HBs concentration of DBS samples (p=0.948). Similarly, elution times were found to have no statistically significant effect on the anti-HBs concentration of DBS samples (p=0.381). However, the main effects analysis revealed that the storage temperature did significantly impact the anti-HBs concentration of DBS samples (p=0.002). This may have been due to the long storage of the samples before testing. A study performed by [11] explained that it is important to store DBS cards appropriately (-80°C) to maintain the integrity of the samples. Our study discovered that the ideal storage temperature for long-term storage of DBS cards was room temperature (25°C) and because some of the cards were stored in other temperatures (-20°C & -80°C), that may have been the reason behind the statistically significant difference of storage temperatures on the anti-HBs concentration of DBS samples (p=0.002).

This DBS elution protocol optimization study is especially significant in settings with limited resources where maintaining optimal laboratory conditions to maintain the integrity of DBS cards could be challenging and when testing for vaccine efficacy in infants as drawing large volumes from infants is not ideal. The practicability of DBS for serological testing is improved by optimizing elution volumes, elution times, and optimal storage temperatures for a reliable DBS elution protocol for anti-HBs quantification. This study will aid in advancing the larger objective of creating an alternative point-of-care diagnostics method for assessing vaccine efficacy and aiding in the effort to eradicate HBV as a public health concern by 2030.

## LIMITATIONS

This study has produced an elution protocol that works for DBS cards stored for long periods. However, due to the lack of sufficient DBS disc for each DBS card stored at the three varying temperatures we had to resort to comparing the results from the DBS samples to their corresponding plasma samples and the study was not able to compare the different factors considered in this elution study to each other. The lack of sufficient DBS discs also led to present results with and without outliers for transparency reasons as the study did not have enough samples to re-test or test in duplicates, specifically samples that were outliers.

## CONCLUSION

To sum up, this study offers insightful information about how to best elute anti-HBs from DBS cards. To improve the reliability of serological testing, it is evident that 450ul of PBS is the ideal elution volume, and agitating DBS discs on a plate shaker for a maximum of 3 hours will yield the best results. The recommendation that 25°C be used as the ideal storage temperature allows DBS cards to be used in locations with limited resources. These results support continued efforts to improve diagnostic accuracy in various healthcare settings by standardizing the use of DBS cards.

## ACKNOWLEDGEMENTS

We thank the CD4 laboratory staff at the Botswana Harvard AIDS Institute Partnership for providing our study with residual plasma and DBS samples.

## POTENTIAL CONFLICTS OF INTEREST

The authors report no conflicts of interest.

## FUNDING

This research was funded in whole, or in part, by the Wellcome [Grant number 218770/Z/19/Z] and SANTHE. For the purpose of open access, the author has applied a CC BY public copyright license to any Author Accepted Manuscript version arising from this submission. MA and BBP were supported by Wellcome. SM was supported by NIH Fogarty International Centre (K43TW012350). SG, WTC and BBP were partially supported by Pan African Bioinformatics Network for the Human Heredity and Health in Africa (H3Africa) consortium (H3ABioNet) and grants from HHS/NIH/National Institute of Allergy and Infectious Diseases (NIAID) (5K24AI131928-04; 5K24AI131924-04) and National Institutes of Health Fogarty International Centre D43TW009610-09S1. H3ABioNet is supported by the National Institutes of Health Common Fund [U41HG006941]. H3ABioNet is an initiative of the Human Health and Heredity in Africa Consortium (H3Africa) program of the African Academy of Science (AAS). SM and BBP were partially supported by the Trials of Excellence in Southern Africa (TESA III) which is part of the EDCTP2 program supported by the European Union (grant number CSA2020NoE3104 TESAIII CSA2020NoE). We also thank the Ministry of Health and Wellness and the Botswana Harvard Health Partnership for the resources they contributed towards the successful completion of this project. The views expressed in this publication are those of the authors and not necessarily those of AAS, NEPAD Agency, Wellcome Trust, EDCTP, BMGF, FIND, or the U.K. government. The funders had no role in the study design, data collection, and decision to publish, or in the preparation of the manuscript.

## Notes

### Competing Interest Statement

The authors have declared no competing interest.

